# Generation of rat lungs by blastocyst complementation in Fgfr2b-deficient mouse model

**DOI:** 10.1101/2022.01.05.475149

**Authors:** Shunsuke Yuri, Yuki Murase, Ayako Isotani

**Author notes:** <u>Corresponding Authors</u>: Shunsuke Yuri and Ayako Isotani, <u>Contact:</u> e-mail address. These authors contributed equally to this work (Shunsuke Yuri, Yuki Murase).

## Abstract

Regenerative medicine is a tool to compensate for the shortage of lungs for transplantation, but it remains difficult to construct a lung *in vitro* due to the complex three-dimensional structures and multiple cell types required. A blastocyst complementation method using interspecies chimeric animals has been attracting attention as a way to create complex organs in animals, but successful lung formation has not yet been achieved. Here, we applied a “reverse-blastocyst complementation method” to clarify the conditions required to form lungs in an Fgfr2b-deficient mouse model. We then successfully formed a rat-derived lung in the mouse model without generating a mouse line by applying a tetraploid-based organ-complementation method. Importantly, rat lung epithelial cells retained their developmental timing even in the mouse body. This result provides useful insights regarding the need to overcome the barrier of species-specific developmental timing in order to generate functional lungs in interspecies chimeras.

## Introduction

The lungs are an interface for gas exchange from oxygen to carbon dioxide through respiration, and are essential for maintaining animal life. Since pulmonary alveoli do not regenerate once damaged, lung conditions such as chronic obstructive pulmonary disease (COPD), the third leading cause of death in the world, are progressive and incurable^1^. Although the only fundamental treatment for COPD or end-stage lung disease is lung transplantation, donor shortage is a critical limitation^2^. To overcome this problem, biological artificial lungs have been created *in vitro* using a decellularized matrix scaffold. Decellularized lungs filled with endogenous lung epithelial cells have been successfully transplanted with a life of a few hours, but are yet to offer a long-term solution^3, 4^. Lung epithelial cells differentiated from human induced pluripotent stem cells (iPSCs) can repopulate into the required scaffold^5^, but to generate a human-scale lung, applying xeno-organs which contain different species as scaffolds has immunological problems^6^.

The lungs develop from epithelial tissue derived from the foregut endoderm, and from mesenchymal tissue derived from the visceral mesoderm. At E9.5, the lung bud bifurcates anteriorly from the ventral foregut endoderm into the mesenchymal tissue^7^. By E16.5, the basic structures of the lung are formed, including airways and terminal bronchi. The epithelium produces basal, ciliated, secretory, and neuroendocrine cells, while the mesenchyme produces smooth muscle, chondrocytes, vascular endothelial cells, and lymphocytes. Next, in E16.5-E17.5, the surrounding mesenchyme become thinner and capillary vessels are actively formed. At this time, type I and II alveolar epithelial cells, and lipofibroblasts arise. In the terminal cyst stage (E17.5-P5), alveolar sacs are formed, surfactant protein production begins, and capillaries develop^8^.

Fibroblast growth factor 10 (Fgf10), which is essential for lung bud elongation, is secreted by the mesenchyme surrounding epithelial tissue^9, 10^, and its signal is accepted by fibroblast growth factor receptor 2 isoform IIIb (Fgfr2b) in the lung epithelium^11^. The interaction of Fgf10 and Fgfr2b is critical for lung development, and both Fgf10 knockout (KO) and Fgfr2b-KO mice showed lung agenesis in previous studies^9, 12^.

To solve the problem of organ shortage, attempts have been made to create donor organs from pluripotent stem cells (PSCs) in the animal body through a process called blastocyst complementation^13-22^. In this method, PSCs such as iPSCs and embryonic stem cells (ESCs) are injected into fertilized eggs of an organ-deficient model. These PSCs can compensate for the defective organs, and PSC-derived organs are created in the body of the resulting chimeric animal. Using this method, transplantable pancreases and thymuses have been successfully produced in interspecies chimeras using mice and rats^13, 14^. PSC-derived kidneys were also generated in interspecies chimeras using mouse PSCs in a rat kidney deficient model, but this process was not successful for rat PSCs in a mouse model^15, 16^. Functional lungs have been produced in intraspecies chimeras using the Fgf10-KO or Fgfr2b-KO mouse models with mouse PSCs, but are yet to be observed in an interspecies chimera^17, 18^. This suggests that the combination of blastocyst and PSC species may be critical, but the exact requirements for successful organogenesis by blastocyst complementation are not yet known. Furthermore, resulting organs are often only partially PSC-derived, even if the model organism exhibits organ-deficient gene dysfunction^15-18^. Therefore, the evaluation of the organ-deficient model is important for generating fully PSC-derived organs and for the realization of future regenerative medicine applications.

This study aimed to examine the organ-deficient model through a “reverse blastocyst complementation method,” which involves the injection of mutant ESCs into wild-type (WT) embryos. The method allows us to efficiently detect mutant cells in the organ and to clarify the conditions for successful lung formation in blastocyst complementation. We achieved lung formation by rat cell complementation in a Fgfr2b-KO mouse model without establishing mouse lines using a tetraploid-based organ-complementation method. The rat cells in the generated lungs unexpectedly retained their developmental timing in the mouse body.

## Results

### The Fgfr2b-KO model was appropriate for organ-complementation of lung epithelium

For a reverse blastocyst complementation system (Fig. 1a), we used mutant ESCs constitutively expressing Su9-DsRed2 (RFP), such that the contribution of mutant cells in the chimera was easily detected. In E14.5 allogeneic chimeric fetuses, RFP-expressing ESC-derived cells were found to have similar contribution rates in various tissues and organs (Supplementary Fig. 1). Therefore, we conducted flow cytometry analysis to estimate the contribution of ESCs to the body tissues of the chimeras based on the percentage of cells that expressed RFP fluorescence.

**Fig. 1.**
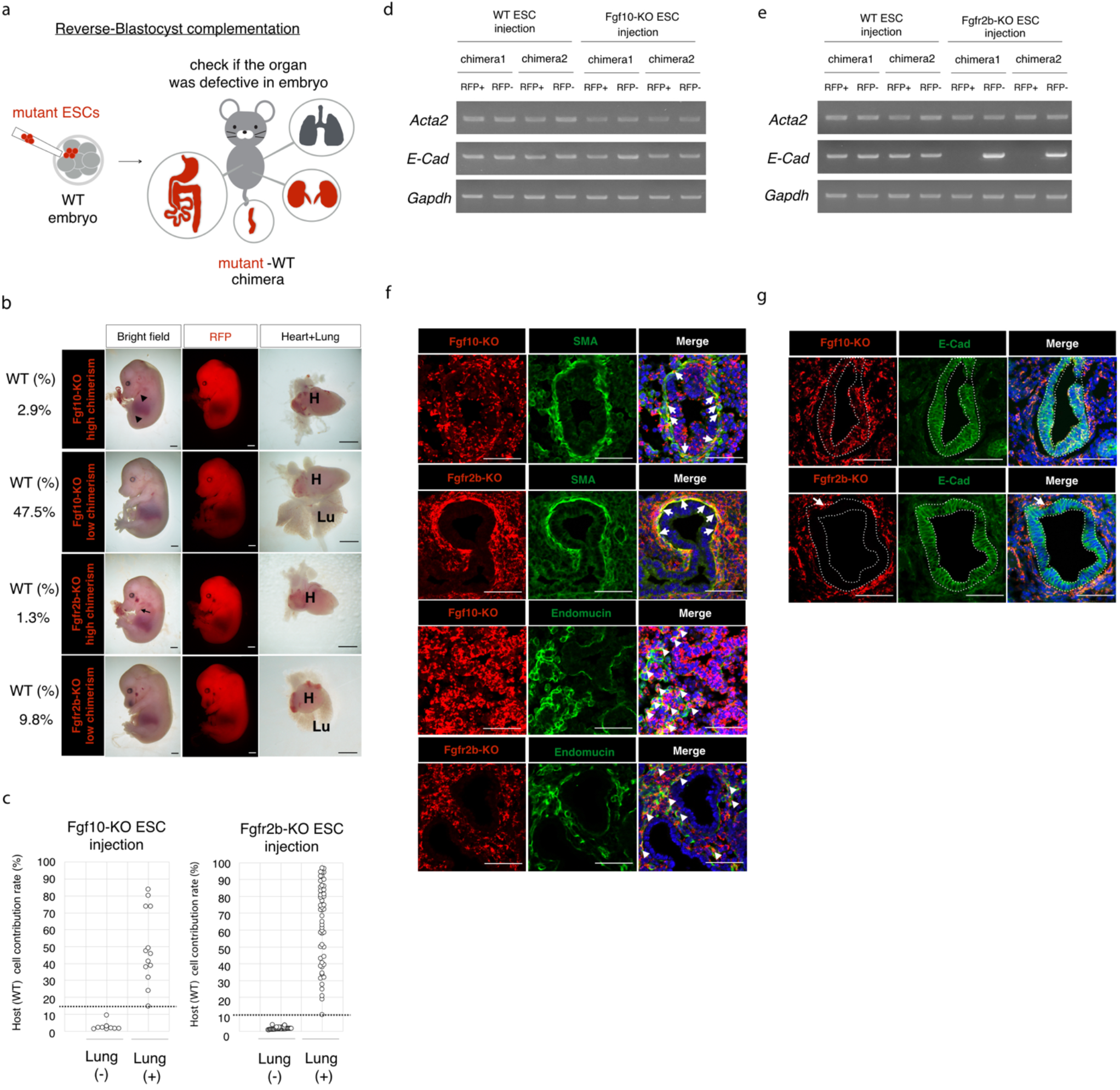
Analysis of Fgf10-KO and Fgfr2b-KO models in lung with reverse-blastocyst complementation. **a** Schematic of reverse-blastocyst complementation. Mutant embryonic stem cells (ESCs) expressing red fluorescent protein (RFP) were injected into the wild type (WT) embryo. Obtained chimeras derived from mutant and WT cells were dissected to determine whether target organ was present. **b** Chimeric embryos derived from Fgf10-knockout (KO) and WT cells or Fgfr2b-KO and WT cells. Chimera with higher contribution of Fgf10-KO cells showed forelimb, hindlimb (black arrowhead), and lung defects. Chimera with higher contribution of Fgfr2b-KO cells showed forelimb (black arrow) and lung defect. H: heart, Lu: lung. Scale bars, 1 mm. **c** Relationship between the cellular contribution rate of the host (WT) cells and the presence of the lung for WT embryos with Fgf10-KO or Fgfr2b-KO ESCs. **d** Gene expressions of *Acta2, E-Cad* or *Gapdh*. RFP+ or RFP-cells were sorted from lungs of Fgf10-KO ESCs and WT cell or WT ESCs and WT cell chimeras. **e** Gene expressions of *Acta2, E-Cad* or *Gapdh*. RFP+ or RFP-cells were sorted from lungs of Fgfr2b-KO ESCs and WT cell or WT ESCs and WT cell chimeras. **f** Immunostaining of SMA, Endomucin in lung of Fgf10-KO and WT cell or Fgfr2b-KO and WT cell chimeras. White arrows or arrowheads indicate that Fgf10-KO or Fgfr2b-KO cells localized at SMA- or Endomucin-positive cells, respectively. Scale bars, 50 µm. **g** Immunostaining of E-Cadherin in lung of Fgf10-KO and WT cell or Fgfr2b-KO and WT cell chimeras. White arrow indicates that Fgfr2b-KO cells localized at E-Cad positive cells. Scale bars, 50 µm.

We designed gRNAs on either side of exon1 on the Fgf10 gene, which contains a start codon, and established the Fgf10-KO ESC lines (Supplementary Fig. 2a, b). We also designed two gRNAs to remove the IgIIIb domain of Fgfr2 and generate Fgfr2b-KO ESC lines (Supplementary Fig. 3a, b). Two ESC lines for Fgf10-KO or Fgfr2b-KO, which have different mutations, were used to produce chimeras (Supplementary Fig. 2c, 3c). Chimeric embryos were generated by injecting Fgf10-KO or Fgfr2b-KO ESCs into E2.5 stage of WT embryos, which were dissected at E14.5 (Table 1, 2). Fgf10-KO chimeras with over 90.7% contribution of Fgf10-KO cells exhibited defects in lung and limb formation (Fig. 1b, c). However, limb and lung defects were not observed in chimeras with more than 14.7% contribution of WT cells (Fig. 1b, c). These results suggest that a certain amount of cells expressing Fgf10 as WT cells is required for lung formation. Similar to the Fgf10-KO chimeras, Fgfr2b-KO chimeras with less than 3.8% WT cell contribution showed defects in the lungs and forelimbs (Fig. 1b, c). Unlike the Fgfr2b-KO phenotype described in a previous study (12), hindlimb defects were not observed in our model. Forelimbs and lungs were observed in chimeras with a WT cell contribution of more than 9.8% (Fig. 1b, c), indicating that proper forelimb and lung development was enabled by 10% contribution of WT cells derived from fertilized eggs.

**Table 1.** Result of Fgf10-KO ESC injection with reverse-blastocyst complementation.

| ESC line | transplantation | implantation | live embryos | chimera |
| --- | --- | --- | --- | --- |
| Fgf10-KO (2E) | 45 | 17 | 17 | 12 |
| Fgf10-KO (7C) | 43 | 24 | 17 | 14 |

**Table 2.** Result of Fgfr2b-KO ESC injection with reverse-blastocyst complementation.

| ESC line | transplantation | implantation | live embryos | chimera |
| --- | --- | --- | --- | --- |
| Fgfr2b-KO (5F) | 180 | 83 | 59 | 44 |
| Fgfr2b-KO (11D) | 166 | 104 | 87 | 75 |

Next, RFP + cells derived from mutant ESCs and RFP – cells derived from WT embryos were sorted from chimeric lungs at E14.5, and the *Acta2* and *E-Cad* expression was examined. *Acta2*, which is expressed in smooth muscle cells that are differentiated from the lung mesenchyme, was detected in the RFP + group of Fgf10-KO and Fgfr2b-KO derived cells (Fig. 1d, e). This indicated that Fgf10-KO and Fgfr2b-KO cells could differentiate into smooth muscle cells. *E-Cad*, which is expressed in lung epithelial cells, was also detected in the RFP + group of Fgf10-KO derived cells but not in that of Fgfr2b-KO derived cells (Fig. 1d, e). This suggests that Fgfr2b-KO cells did not contribute to the lung epithelium. To further investigate, sections of Fgf10-KO and WT chimeric lungs were immunostained for mesenchymal cell-derived tissues, such as smooth muscle or vasculature with smooth muscle actin (SMA), or for endomucin antibody, respectively. Fgf10-KO and Fgfr2b-KO cells contributed to the smooth muscle and vasculature cells (Fig. 1f). We also immunostained the epithelial tissues using antibodies against E-cadherin. Fgf10-KO cells contributed to the lung epithelium (number of lung epithelial ducts without Fgf10-KO cells = 0/16; Fig. 1g). These results indicated that the Fgf10-KO animal model did not provide any organ niches in the lungs for WT cell compensation. In contrast, Fgfr2b-KO cells did not contribute greatly to lung epithelial cells (number of lung epithelial tubules without Fgfr2b-KO cells = 77/81). However, even in the four cases of lung epithelial ducts to which Fgfr2b-KO cells contributed, only a few Fgfr2b-KO cells were identified (Fig. 1g). These results show that the Fgfr2b-KO animal model provided an organ niche for lung epithelial tissues, and only 10% contribution of WT cells was necessary for proper lung formation.

### Mouse ESCs promoted lung formation in Fgfr2b-KO mouse model via a tetraploid-based organ-complementation method

Since conventional blastocyst complementation method is time-consuming, we next generated mutant and WT chimeras from two types of ESCs (mouse Fgfr2b-KO and WT) using the tetraploid-based organ-complementation method^23^. We first injected RFP-expressing mouse Fgfr2b-KO ESCs into tetraploid embryos at the E2.5 stage, followed by GFP-expressing mouse WT (G-mWT) ESCs injected at the E3.5 stage (Fig. 2a). After transplantation of the chimeric embryos, we obtained E14.5 chimeric fetuses (n=2; Table 3). Similar to the chimeras obtained using the reverse-blastocyst complementation method, defects in the forelimbs and lung epithelium were complemented by cells derived from G-mWT-ESCs (Fig. 2b). We examined whether the lung epithelium was complemented with G-mWT-ESCs by immunostaining with an E-cadherin antibody. In the chimeric lungs, the epithelial tissue was found to be complemented by GFP-expressing cells (n=12/12) (Fig. 2c). To address whether the resulting lung was functional after birth, we performed a caesarean section at E19.5, and obtained two chimeras (Table 3). Both chimeras showed a normal appearance, but the non-chimeras that did without GFP signal showed cyanotic skin color (Fig. 2d). Although lungs were not present in the non-chimera, as expected, GFP-expressing lungs were present in chimeras (Fig. 2e, Supplementary Fig. S4). This indicates that the mouse WT ESCs were able to correct the lung epithelium defect in the Fgfr2b-KO model through tetraploid-based organ complementation.

**Table 3.** Result of Fgfr2b-KO ESC and G-mWT ESC injection with tetraploid complementation at E14.5 and P0.

| Stage at dissection | transplantation | implantation | live embryos | chimera |
| --- | --- | --- | --- | --- |
| E14.5 | 48 | 18 | 4 | 2 |
| P0 | 27 | 5 | 2 | 2 |

**Fig. 2.**
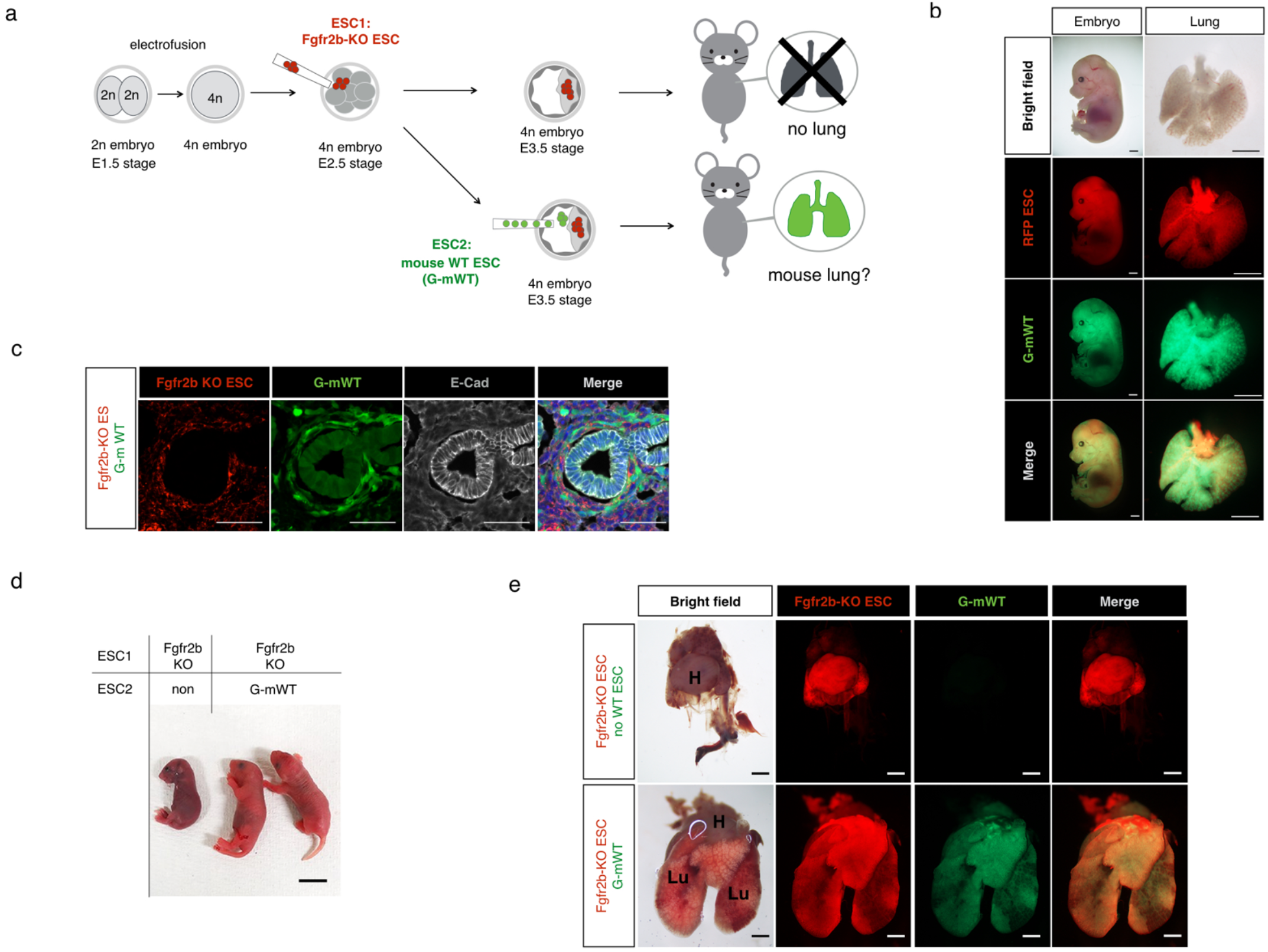
Generation of mouse ESC-derived lung in Fgfr2b-KO model with tetraploid-based organ complementation method. **a** Schematic for producing chimeras from Fgfr2b-knockout (KO) and mouse wild type (WT) ESCs. Two cell-stage embryos were electrically fused to produce a 4n embryo. Fgfr2b-KO ESCs were injected at the E2.5 stage, followed by GFP-expressing mouse WT (G-mWT) ESCs at the E3.5 stage. Without G-mWT ESC injection, lung agenesis was theoretically observed (upper panel), then examined to determine whether G-mWT ESCs overcame lung agenesis (lower panel). **b** Embryo and lung derived from red fluorescent protein (RFP)-expressing Fgfr2b-KO ESCs and G-mWT ESCs chimera. Scale bars, 500 µm. **c** Immunostaining of E-Cad in lung derived from Fgfr2b-KO ESCs and G-mWT ESCs chimera. Note that lung epithelial cells were composed by GFP-expressing mouse WT cells. Scale bars, 50 µm. **d** Neonates from obtained chimeras. No WT ESC-contributed pups showed cyanosis. Scale bars, 1 cm. **e** GFP and RFP images of isolated heart and lungs from Fgfr2b-KO only and Fgfr2b-KO and G-mWT ESC chimeras at P0. (H: heart, Lu: lung) Scale bars, 1 mm.

### Rat ESCs promoted lung formation in Fgfr2b-KO mouse model using a tetraploid-based organ-complementation method

Since the mouse WT ESCs ameliorated the lung epithelial defect in Fgfr2b-KO mice, we next examined whether rat lungs could be formed in the Fgfr2b-KO lung deficient mouse model using tetraploid-based organ-complementation. When GFP-expressing rat WT (G-rWT) ESCs were injected into WT mouse fertilized eggs, rat cells contributed unevenly to the embryos depending on the organ in the resulting a mouse-rat interspecies chimera. Notably, rat cells contributed more to the lungs than other to organs, whereas G-mWT ESC-derived cells uniformly contributed to each organ^24^ (Supplementary Fig. 5a-5c). Therefore, we hypothesized that the lung epithelial defect of mouse Fgfr2b-KO mice could be complemented by rat cells. We injected RFP-expressing mouse Fgfr2b-KO ESCs into tetraploid embryos at the E2.5 stage, followed by G-rWT-ESCs injected at the E3.5 stage (Fig. 3a). At E14.5, we observed successful lung formation in the interspecies chimeric fetuses (Fig. 3b) (n=3; Table 4). This indicated that the defective lung phenotypes from Fgfr2b-KO mice could be recovered with rat cells, although the obtained lungs were relatively small compared to those resulting from mouse WT cells (Fig. 3b, Supplementary Fig 6a vs Fig. 2b). We also confirmed that most of the tubular structures that appeared to be epithelial tissue in interspecies chimeric lungs were complemented by rat cells (n= 12/13) (Fig. 3c and Supplementary Fig. 6b). To examine whether the lung rescued by rat WT cells was functional after birth, we performed a caesarean section at E19.5, and obtained two chimeras (Table 4). However, both pups with GFP fluorescence (rat chimera) showed cyanotic skin color similar to the pups that did not show GFP signal (non-chimera) (Fig. 3d). The non-chimera pups died within 21 min, on average, due to respiratory failure, whereas the Fgfr2b-KO chimeras with G-mWT cells survived for more than 5 h (Fig. 3e). The rat chimeras showed postnatal mortality within 10 min to 15 min (Fig. 3e), even though exhibited GFP-expressing lungs (Fig. 3f and Supplementary Fig. 7). These results showed that rat WT ESCs did promote lung epithelial development in the Fgfr2b-KO mouse model with tetraploid-based organ-complementation method, but the generated lungs were not functional after birth.

**Table 4.**
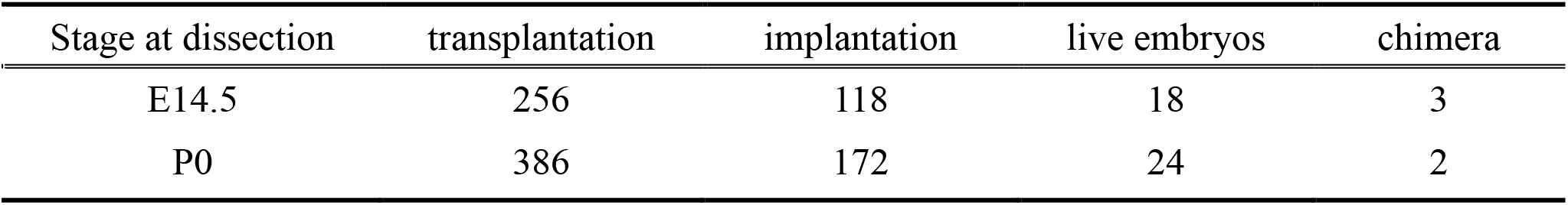
Result of Fgfr2b-KO ESC and G-rWT ESC injection with tetraploid complementation at E14.5 and P0.

| Stage at dissection | transplantation | implantation | live embryos | chimera |
| --- | --- | --- | --- | --- |
| E14.5 | 256 | 118 | 18 | 3 |
| P0 | 386 | 172 | 24 | 2 |

**Fig. 3.**
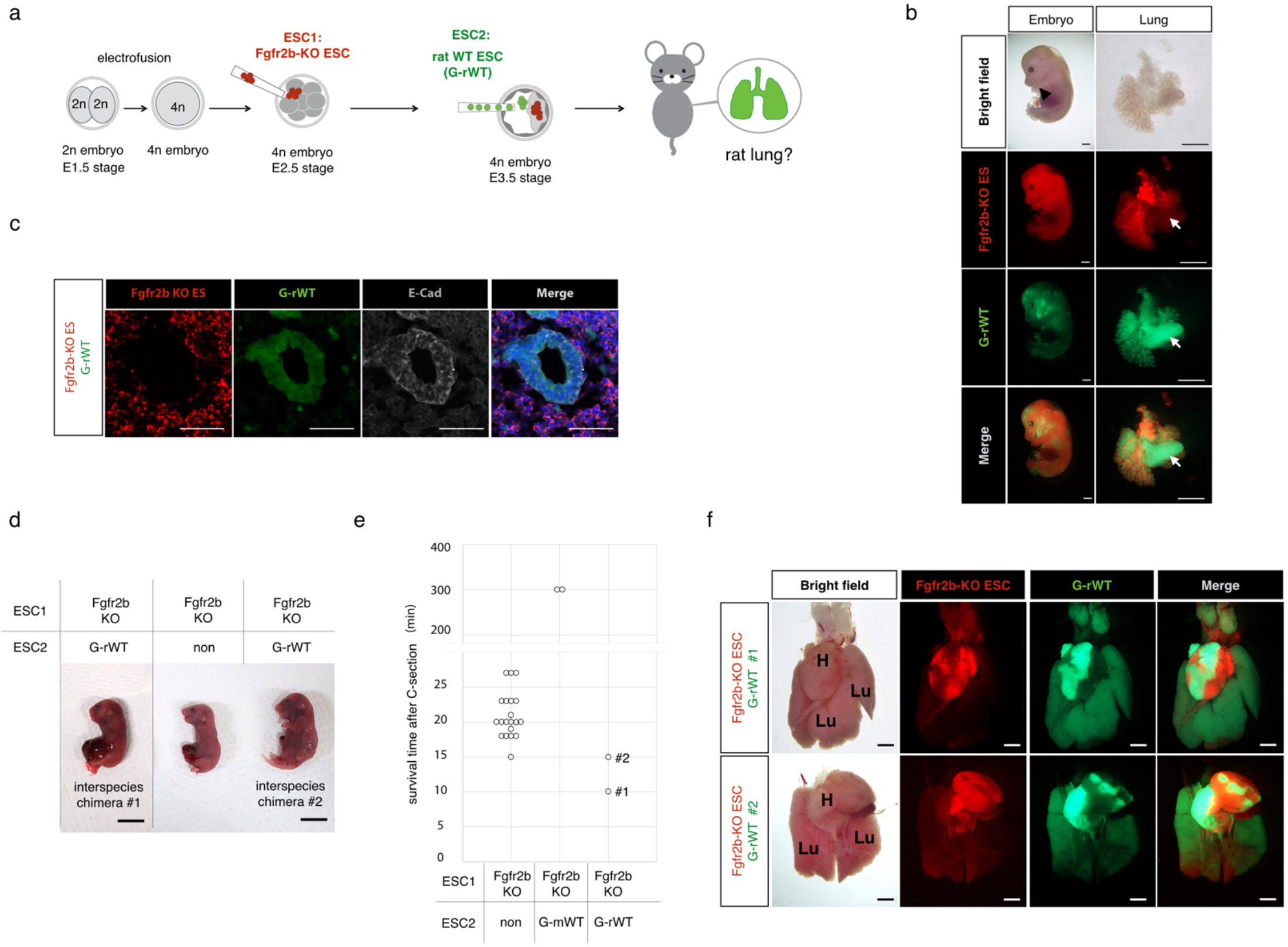
Generation of rat ESC-derived lungs in Fgfr2b-KO model with tetraploid-based organ complementation method. **a** Schematic for producing chimeras from Fgfr2b-knockout (KO) ESCs and rat wild type (WT) ESCs. Two cell-stage embryos were electrically fused to produce a 4n embryo. Fgfr2b-KO ESCs were injected at the E2.5 stage, followed by GFP-expressing rat WT (G-rWT) ESCs at the E3.5 stage. **b** Embryo and lung derived from Fgfr2b-KO and G-rWT ESC chimera at E14.5. Chimera with G-rWT ESCs have forelimb (black arrowhead) and lung. Note that one of the lung lobes was almost fully composed of rat cells (white arrow). Scale bars, 500 µm. **c** Immunostaining of E-Cad in lung derived from Fgfr2b-KO and G-rWT ESCs chimera. Note that lung epithelial cells were composed of G-rWT cells. Scale bars, 50 µm **d** Neonates from obtained chimeras. No WT ESC-contributed pups or G-rWT ESC chimeras showed cyanosis. Scale bars, 1 cm **e** Survival time of obtained pups after Caesarian section (C-section) (min). **f** GFP and RFP images of isolated heart and lungs from Fgfr2b-KO and G-rWT ESC chimeras (#1, #2) at P0. Note that lung from Fgfr2b-KO and rat ESCs #1 was composed of almost all rat cells. (H: heart, Lu: lung) Scale bars, 1 mm.

### Lung epithelium formed by rat ESCs preserved intrinsic developmental time in the Fgfr2b-KO mouse model

Since the morphology of the lungs in the rat chimera was not the same as that in the mouse chimera for the Fgfr2b-KO model (Fig. 2e vs Fig. 3f), we further analyzed the generated lungs. Histology of lungs composed of Fgfr2b-KO and G-mWT cells revealed normal saccular expansion and septal thinning, similar to that of the mouse WT control, suggesting that the G-mWT-ESCs could fully compensate for the lung dysfunction in the Fgfr2b-KO model. In contrast, histological analysis revealed that lungs from Fgfr2b-KO and G-rWT cells showed abnormal alveolar expansion with smaller airspaces and much smaller alveoli, indicating that lung development was delayed compared to the intraspecies model, or that no surfactant protein was secreted and the lungs failed to inflate with air (Fig. 4a, b). To examine whether the histological abnormality of the lung from Fgfr2b-KO and G-rWT chimeras was from dissection timing, we dissected mouse WT control or mouse WT and G-rWT interspecies chimeras at 10 min after caesarean section. However, we did not observe a difference in air space between the WT mouse and the G-rWT interspecies chimera (Fig. 4a). This suggested that the abnormality of the lung from Fgfr2b-KO and G-rWT chimeras was not due to dissection timing. Since rat development is slower than mouse development, we further investigated the immaturity of the lungs of the Fgfr2b-KO and rWT chimeras. We immunostained for the SRY-box containing gene 9 (Sox9), a marker of lung distal epithelial progenitor cells. Sox9 is mainly expressed in the epithelial progenitor cells from E11.5–E16.5, and is hardly detectable by E18.5^25, 26^. We could not detect Sox9 positive progenitor cells in the lungs of Fgfr2b-KO and G-mWT chimeras (Fig. 4c). However, in the lungs of Fgfr2b-KO and G-rWT chimeras, numerous Sox9 positive progenitor cells were detected in the epithelial tubules (Fig. 4c). We also could not detect Sox9 positive progenitor cells in the lungs of the mouse WT control or the mouse WT and G-rWT interspecies chimera dissected 10 min after caesarean section, even though the GFP-expressing rat cells were highly populated in the lungs (Supplementary Fig. 8a, b). In the lungs of mouse WT and G-rWT interspecies chimeras, we found that GFP-expressing rat cells were almost not observed in the lung epithelium (Supplementary Fig. 8b). *Sftpc* mRNA, which is expressed in mature AT2 differentiated from lung epithelial progenitor, was not detected or present in low levels in some lung lobes of the mouse WT and G-rWT interspecies chimeras (Supplementary Fig. 8c, d). These results indicated that rat cells were likely to be eliminated from lung epithelial tissue when mouse WT cells were present. Together, the Fgfr2b-KO and rat WT chimeras could not breathe after birth, likely because rat lung epithelial cells preserved their own developmental timing even in the mouse body.

**Fig. 4.**
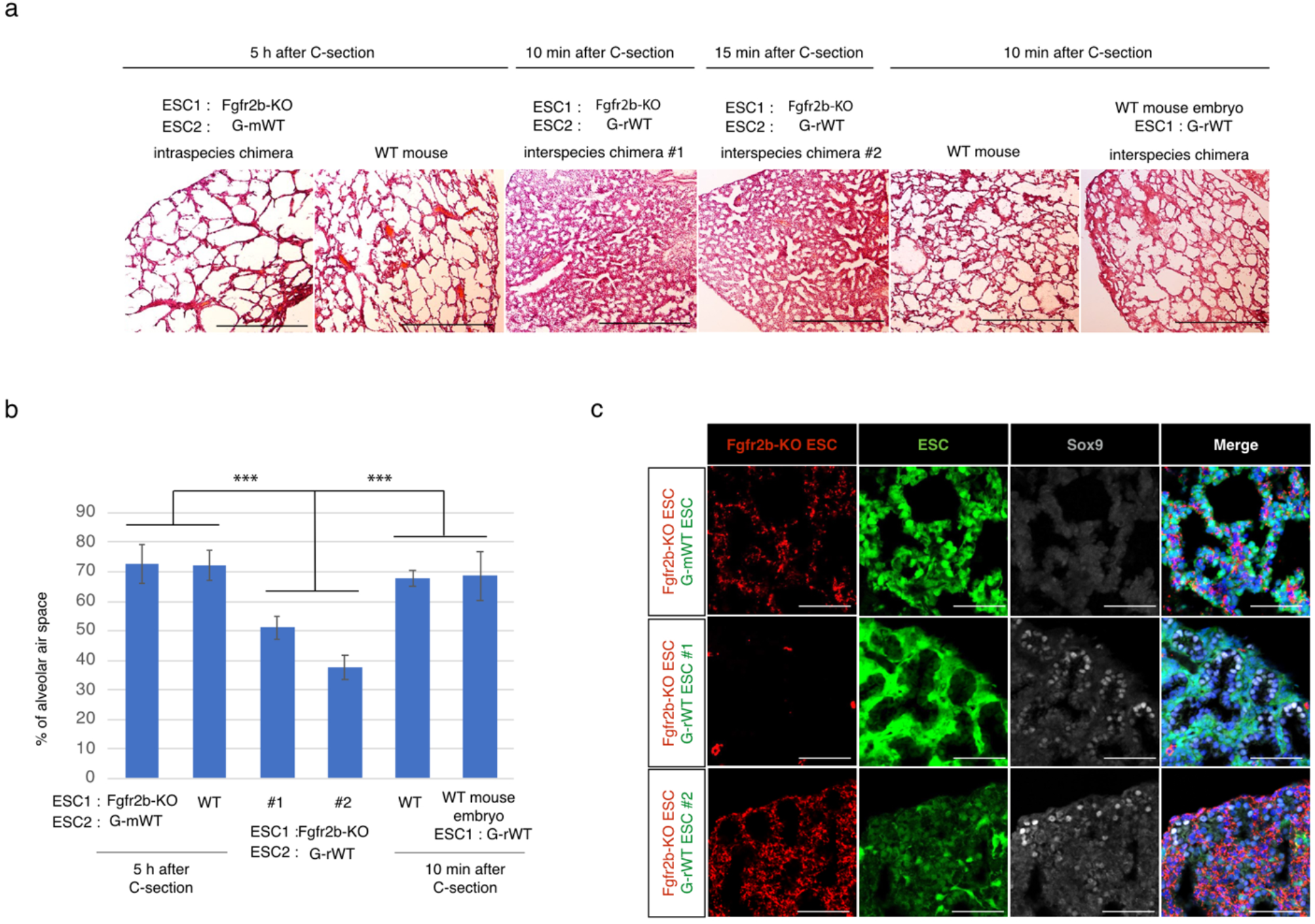
Analysis of rat-derived lung with Fgfr2b-KO model in neonatal stage. **a** Hematoxylin and Eosin (H&E) stain of the lung sections at P0. Lungs from Fgfr2b-knockout (KO) ESC and GFP-expressing mouse WT (G-mWT) ESC chimeras or WT mice were dissected 5 h after Caesarian section (C-section). Lungs from Fgfr2b-KO ESC and GFP-expressing rat WT (G-rWT) ESC chimeras (#1, #2) were dissected after chimeras died (#1: 10 min, #2: 15 min). Lungs from WT mice or WT mouse and G-rWT ESC chimeras were dissected 10 min after C-section. Scale bars, 500 µm **b** The air space was measured from the obtained lung section in Fig.4a. Numbers indicate the percentage of alveolar air space. (*n*=3 for each). Data are presented as means ± S.D. from three independent regions. ***: *p*<0.01 **c** Immunostaining of Sox9 (Gray) in the lungs derived from Fgfr2b-KO ESCs (Red) and G-mWT or G-rWT ESC (Green) chimeras. Scale bars, 100 µm

## Discussion

The production of organs by blastocyst complementation has been highlighted as one of the most promising regenerative medicine platforms. One of the remaining problems is the lack of knowledge regarding successful production of PSC-derived organs. In this study, we applied a reverse-blastocyst complementation method to evaluate an organ-deficient model for the production of lungs by blastocyst complementation, establishing an important benchmark that will allow us to evaluate whether a given organ-deficient model can provide an appropriate organ niche. Consequently, we clarified the success conditions for generating lungs through the blastocyst complementation method and achieved the production of lungs with rat cells in a mouse lung-deficient model.

Here, we demonstrate the effectiveness of the reverse-blastocyst complementation method. Reverse-blastocyst complementation has been applied in previous studies to analyze whether the abnormalities caused by gene knockout in mutant embryos are intrinsic defects due to their gene functions (cell-autonomous) or extrinsic defects due to the surrounding microenvironment (cell non-autonomous)^27-30^. Compared to the blastocyst complementation method, the reverse-blastocyst complementation method provides several advantages for evaluating organ-deficient animals. First, the analysis of organ deficient-WT chimera embryos in the blastocyst complementation method was time-consuming and inefficient because it required the establishment of a genetically modified heterozygous mouse line, as well crossing to obtain even single gene deficient embryos, which appears in only one in four. In the reverse-blastocyst complementation method, only mutant PSCs need to be established to analyze organ deficient-WT chimeras, and organ deficient-WT chimeras can be obtained from all embryos injected with mutant PSCs. In addition, using CRISPR-Cas9 technology, it is possible to obtain mutant PSCs with high efficiency^31^. Second, in blastocyst complementation, if the gene plays a pivotal role in extraembryonic tissue, injected PSCs cannot compensate for the abnormality. Since PSCs only differentiate into a given embryonic lineage, the abnormality from gene mutation occurs only in the embryonic tissue in reverse-blastocyst complementation. Third, in the blastocyst complementation method, injected PSCs usually express fluorescent proteins^17,18^, so their distribution in the tissue can be easily tracked. However, if the contribution of PSC-derived cells is high, it is difficult to determine the contribution of mutant-derived cells in the tissue^15-18^. In contrast, with the reverse-blastocyst complementation method, the distribution of mutant PSC-derived cells in the tissue can be readily ascertained, even if the mutant cell contribution is low.

We investigated the lungs of Fgf10-KO or Fgfr2b-KO and WT chimeras using the reverse-blastocyst complementation method. Recently, Kitahara et al.^17^ generated lungs in an Fgf10-KO mouse model using blastocyst complementation. Consistent with our results, they showed that Fgf10-KO cells were included in most cell types in the lungs. However, they observed that WT PSC-derived cells were primarily detected in the epithelial cells of the generated lungs, which we did not observe. Since Fgf10 signaling regulates differentiation into lipofibroblasts in the lung mesenchyme, which appear from E15.5^32^, Fgf10-KO cells may be able to change their distribution in the lungs after the E15.5 stage. Mori et al.^18^ also showed that mouse PSCs could generate functional lungs in an Fgfr2 conditional KO model through the blastocyst complementation method, and the epithelial tissues of generated lungs were highly populated by PSC-derived cells compared to lung mesenchyme tissue. Consistent with their results, we observed that Fgfr2b-KO cells could not contribute to the lung epithelium through the reverse-blastocyst complementation method. In summary, we were able to demonstrate the feasibility of the reverse-blastocyst complementation method to evaluate the contribution of WT cells to organs without generating organ-deficient mice.

Since we realized that mouse WT cells contributed fairly evenly to all tissues in E14.5 embryos, we could roughly estimate chimerism in the target organs of the organ-deficient model. Based on this estimation, the presence of a certain number of normal cells was necessary to overcome the phenotype of lung agenesis in Fgf10-KO or Fgfr2b-KO models. If the presence of a certain number of normal cells is also important for the formation of kidney, then rat cells might not be able to rescue the mouse kidney agenesis model^16^, because we realized that rat cells cannot contribute much to the mouse kidney in this study. In the future, we will need insight into the percentage of WT cells required to produce the organs of each tissue using the reverse-blastocyst complementation method.

We applied a tetraploid-based organ-complementation method, which also allows the production of WT-PSC -and KO-PSC-derived chimeras without generating mouse lines, to produce rat lungs in the Fgfr2b-KO mouse model. While higher WT contribution in the defective organ is expected to be one of the critical factors when interspecies chimeras are produced for organ generation^33^, we and others^24^ found that rat PSCs were likely to contribute to mouse lung tissue. In addition, we realized that few WT-derived cells were required to prevent lung agenesis in the Fgfr2b-KO model in this study. Indeed, we succeeded in overcoming the lung defect in Fgfr2-KO mice with rat PSCs through the tetraploid complementation method. The size of all three lungs complemented by rat PSCs was smaller than that of lungs complemented by mouse PSCs at E14.5. In particular, the lung size was smallest in Fgfr2b-KO and rat WT chimeras, with the lowest percent contribution from rat cells. Since lung size is determined by the number of lung epithelial progenitor cells^34^, this may indicate that lung size is also determined by the number of rat lung epithelial progenitor cells in interspecies chimera.

Intriguingly, in one of the resulting rat lungs at postnatal stage (Fig. 3f, chimera #1), the rat epithelial progenitor cells remained immature, even if the lung was composed almost entirely of rat cells. This result suggests that rat lung epithelial cells retain their own developmental speed despite their presence in a different species, or that rat lung mesenchymal cells, including lung epithelial cells, are delayed compared to mouse development. In another example (Fig. 3f, chimera #2), rat lung epithelial cells were still in an immature state, even though most of the mesenchymal cells were mouse cells. This result raises the possibility that rat epithelial cells are unable to receive signals from mouse mesenchymal cells. Similar to this observation, rat germ cells, even when present in mouse testes, have been shown to differentiate at the time typical for rats and therefore generate the structural pattern of rat spermatogenesis^35^. This suggests that some cells exhibit intrinsic regulation of differentiation, even in interspecies environments. Therefore, the generation of functional lungs in interspecies chimeras may require overcoming the barrier of species-specific intrinsic developmental timing.

There are also some limitations in our study. Although rat lungs were formed in the Fgfr2b-KO mouse model, they were not functional after birth because the rat lung epithelial tissue was still in the prenatal stage. Balancing the proliferation and differentiation of lung epithelial progenitor cells requires fine control of Sox9 expression levels^36^. Thus, modulation of Sox9 may be important for generating functional lung in interspecies blastocyst complementation. Sox9 expression is regulated by Nmyc or Asxl1 ^26, 37^. Therefore, it may be important to modulate the expression of Nmyc or Asxl1 to regulate the intrinsic developmental timing of lung epithelial progenitor cells. In addition to Sox9, Creb1, Grhl2, Carm1, and Foxm1 are also required for appropriate alveolar formation and development during fetal lung development^38-41^. In the future, it may be necessary to determine the factors which is related to the intrinsic developmental timing of lung epithelial progenitor cells in xenogeneic lung. Moreover, future studies comparing species with different developmental speeds other than the mouse-rat combination would be useful to better understand species-specific developmental time.

In summary, our analysis provides evidence that the Fgfr2b-KO demonstrate lung epithelial deficient model by reverse-blastocyst complementation method. With the model, we propose that rat ESCs potentiate the ability to rescue the lung agenesis in mouse. Furthermore, our findings point to regard for species-specific developmental timing is a key point for generating functional lung in interspecies blastocyst complementation.

## Methods

### Animals

All animal experiments were conducted in accordance with the guidelines of “Regulations and By-Laws of Animal Experimentation at the Nara Institute for Science and Technology” and were approved by the Animal Experimental Committee at the Nara Institute of Science and Technology (approval nos. 1639 and 2109). The animal experiments in this study were performed in compliance with ARRIVE guidelines^42^. ICR mice were purchased from Japan SLC, Inc.

### Collection of eggs

ICR females aged 8-10 weeks were treated with CARD HyperOva (Kyudo) and hCG (ASKA Animal Health) for superovulation, then were mated with ICR male mice. Two-cell stage zygotes were collected from female oviducts 42–46 h after hCG injection using the flush-out method. The collected two-cell stage embryos were incubated in KSOM medium at 37 °C under 5% CO_2_ conditions until use.

### Construction of the plasmid vector and design of sgRNA

Oligo DNAs of the target sequence were ligated into the BbsI site of the pSpCas9(BB)-2A-Puro (pX459) V2.0 plasmids (Addgene #62988). The combination of #1 and #2 or #3 and #4 oligos was used to establish Fgf10-KO ESCs, and the combination of #8 and #9 or #10 and #11 oligos was used to establish Fgfr2b-KO ESCs (Supplementary Table1). All oligonucleotides were designed using the CrisperDirect website to identify specific target sites.

### Establishment of Fgf10-KO and Fgfr2b-KO ESC lines

Fgf10 or Fgfr2b-KO ESCs were established using the R01-09 ESC line, which was newly established from 129X1 and R01 F1 embryos. R01 mouse lines were established through the tetraploid complementation method from R01 ESCs obtained from Dr. Masahito Ikawa, established from 129 and BDF1 F1 embryos and constitutively expressing RFP by the CAG-Su9-DsRed2 transgene, which localizes to the mitochondria. R01-09 ESCs were seeded on mouse embryonic fibroblasts (MEFs) and then transfected with the two designed plasmids using Lipofectamine 3000 (Thermo Fisher Scientific). Transfected cells were selected by transient treatment with 1 µg/ml puromycin for 2 days, and then ESC colonies were subjected to genotyping with PCR and sequencing. ESCs were cultured on gelatin-coated dishes, and MEFs with N2B27 medium supplemented with 3 µM CHIR99021(Axon; 1386), 1.5 µM CGP77675 (Sigma; SML0314), and mouse LIF (leukemia inhibitory factor) (N2B27-a2i/L medium)^43^.

### Genotyping

Genotyping primers for detecting Fgf10-KO and Fgfr2b-KO ESCs are shown in Table S1. DNA fragments were amplified using GoTaq (Promega) for 40 cycles to detect null or WT alleles under the following conditions: 94°C for 30 s, 60°C for 30 s and 72°C for 60 s.

### Flow cytometry analysis and fluorescence-assisted cell sorting

All chimeric embryos were recovered at the E14.5 stage. Tail, kidney, lung, stomach, and intestine samples were incubated with 0.25% trypsin for 10 min at 37°C. After pipetting to dissociate the tissue, 10% FBS in PBS was added, and samples were filtered through a 37 µm mesh. The FL3 detector on Accuri (BD Bioscience) was used to detect RFP+ populations. Tail samples were used to estimate chimerim on Fgf10-KO or Fgfr2b-KO chimera. SH800SFP (SONY) was used to sort between the RFP+ and RFP-subpopulations.

### RNA expression analysis

Total RNA was purified using Trizol reagent (Thermo Fisher Scientific) and used for reverse transcription. cDNA was prepared using the SuperScript IV VILO master mix (Thermo Fisher Scientific). RT (reverse transcription) PCR was performed using GoTaq (Promega). For quantitative RT-PCR analysis, Luna Universal qPCR Master Mix (NEB) was used to amplify the DNA fragment, and amplified DNA was detected on a LightCycler 96 (Roche). The primers used for RT-PCR are described in Supplementary Table2.

### Tetraploid complementation and rat ESC injection

Tetraploid embryos were prepared as described previously^44, 45^. In brief, ICR two-cell stage embryos were placed in fusion buffer, and electrofusion was performed using CFB16-HB and LF501PT1-10 electrodes (BEXCo.Ltd,). Tetraploid embryos were incubated in KSOM at 37°C under 5% CO_2_ until use. 6-8 cells of Fgfr2b-KO ESCs were injected into tetraploid embryos at E2.5. These embryos were cultured to the E3.5 stage and injected into 6-8 cells of GFP-expressing rat ESCs (rG104)^46^, followed by transfer into the uterus of E2.5 pseudopregnant ICR mice. The fetuses were recovered and dissected at E14.5, and offspring were recovered at E19.5 via Caesarean section. The ESC-derived offspring were analysis using RFP or GFP signal under fluorescence stereo microscope (MZ FLIII; Leica).

### Immunocytochemistry and HE staining

The lungs at E14.5 or E19.5 were fixed with 4% paraformaldehyde (PFA) in phosphate buffered saline (PBS) (-) (Nacalai) for 15 min at 25 °C or overnight at 4 °C. After washing with PBS, the lungs were immersed in 10, 20, and 30% sucrose. The treated lungs were then sunk into Tissue-TeK O.C.T compound (Sakura Finetek). After making sections with a cryostat (NX70; Leica) at 10 µm, the slides were dried at 25 °C, followed by washing with PBS (-). Slides were immersed in EtOH and 4% PFA solution (1:1) for 15 s and washed with ddH_2_O, then treated with Mayer’s hematoxylin solution (WAKO) for 1 min and washed with PBS (-) and ddH_2_O. Next, slides were immersed in 0.5% eosin solution (WAKO) for 1 min and washed three times with 100% EtOH, then twice more in xylene. Mountquick (DAIDO SANGYO) was added to the slides and samples were covered with cover glass. The sections were observed under a microscope with 10x objective lens (BX60; Olympus). For immunostaining, slides were treated with 1% bovine serum albumin (BSA) (Sigma) for 60 min at 25 °C. The primary antibody was incubated overnight at 4 °C. The slides were then washed three times for 5 min with PBS (-) and incubated with the secondary antibody for 1 h at 25 °C. After washing three times for 5 min with PBS (-) at 25 °C, the nuclei were stained with Hoechst33342 (Dojindo, KV072) and diluted to 1:1000 in PBS for 30 min at 25 °C before a final PBS (-) wash. The antibodies used included: rabbit anti-E-Cadherin (Cell Signaling; #3195), rat anti-E-Cadherin (TAKARA; M110), rat anti-Endomucin (Santa Cruz; sc-65495), mouse anti-SMA (Biolegend; 904601), rabbit anti-Sox9 (Millipore; AB5535), goat Alexa Fluor 488 anti-rabbit IgG (Thermo Fisher Scientific; A11017), goat Alexa Fluor 647 anti-rabbit IgG (Thermo Fisher Scientific; A21246), goat Alexa Fluor 488 anti-mouse IgG (Thermo Fisher Scientific; A11017), goat Alexa Fluor 647 anti-mouse IgG (Thermo Fisher Scientific; A21237), goat Alexa488 Fluor anti-rat IgG (Thermo Fisher Scientific; A11006), and goat Alexa647 Fluor anti-rat IgG (Thermo Fisher Scientific; A21247). Immunostained samples were examined using a laser confocal microscope (LSM700, LSM710, LSM980; Zeiss).

### Air space measurement

The air area fraction at E19.5 was measured from lung sections stained with hematoxylin and eosin. More than three non-overlapping fields (x10 objective lens) from each lung sample were analyzed. The percentage of air space in the total distal lung area was analyzed using the ImageJ software.

### Statistical analysis

For air space measurement, all values are expressed as mean ± standard deviation from at least three different regions in each sample. For quantitative RT-PCR data expressed as relative fold changes, all values are expressed as mean ± standard deviation from at least triplicate experiments. Student’s *t*-test for unpaired comparisons was performed and differences were considered significant when *p* < 0.01.

## Acknowledgements

We thank Dr. Ikawa (Osaka University) for kindly providing R01 ESCs. We also would like to thank Editage (www.editage.com) for English language editing. This work was supported by JSPS KAKENHI Grant Number (16K07091 and 18H04885) to A.I. and (17H06868 and 18K06031) to S.Y., Start Up Fund for female researchers in NAIST to A.I., KAC 40th Anniversary Research Grant to A.I., The NOVARTIS Foundation (Japan) for the Promotion of Science to A.I., Next Generation Interdisciplinary Research Project to A.I., and The foundation for Nara Institute of Science and Technology to S.Y.

## Author contributions

S.Y. and A.I. designed the study. S.Y. wrote the manuscript with help of A.I. S.Y. and Y.M. performed embryo manipulation, analyzed the chimeras, performed molecular biological analysis and cell culture. S.Y. and A.I supervised the project.

## Competing interests

The authors declare that they have no competing interests.

### Additional information

Correspondence and requests for materials should be addressed to S.Y. or A.I.

All data needed to evaluate the conclusions in the paper are presented in the paper and/or the Supplementary information.

## References

1. WHO data on 21 June 2021. https://www.who.int/news-room/fact-sheets/detail/chronic-obstructive-pulmonary-disease-(copd)

2. Valapour, M. et al. OPTN/SRTR 2019 Annual Data Report: Lung. Am. J. Transplant. 21, 441–520 (2021).

3. Ott, H. C. et al. Regeneration and orthotopic transplantation of a bioartificial lung. Nat. Med. 16, 927–933 (2010).

4. Petersen, T. H. et al. Tissue-Engineered Lungs for in Vivo Implantation. Science 329, 538–541 (2010).

5. Ghaedi, M. et al. Bioengineered lungs generated from human iPSCs-derived epithelial cells on native extracellular matrix. J. Tissue Eng. Regen. Med. 12, e1623–e1635 (2018).

6. Stahl, E. C. et al. Evaluation of the host immune response to decellularized lung scaffolds derived from alpha-Gal knockout pigs in a non-human primate model. Biomaterials 187, 93–104 (2018).

7. Herriges, M. & Morrisey. E. E. Lung development: orchestrating the generation and regeneration of a complex organ. Development 141, 502–513 (2014).

8. Loering, S., Cameron, G. J. M., Starkey, M. R. & Hansbro, P. M. Lung development and emerging roles for type 2 immunity. J. Pathol. 247, 686–696 (2019).

9. Sekine, K. et al. Fgf10 is essential for limb and lung formation. Nat. Genet. 21, 138–141 (1999).

10. Yuan, T., Volckaert, T., Chanda, D., Thannickal, V. J. & De Langhe, S. P. Fgf10 Signaling in Lung Development, Homeostasis, Disease, and Repair After Injury. Front. Genet. 9, 418 (2018)

11. Ohuchi, H. et al. FGF10 acts as a major ligand for FGF receptor 2 IIIb in mouse multi-organ development. Biochem. Biophys. Res. Commun. 277, 643–649 (2000).

12. De Moerlooze, L. et al. An important role for the IIIb isoform of fibroblast growth factor receptor 2 (FGFR2) in mesenchymal-epithelial signalling during mouse organogenesis. Development 127, 483–492 (2000).

13. Isotani, A., Hatayama, H., Kaseda, K., Ikawa, M. & Okabe, M. Formation of a thymus from rat ES cells in xenogeneic nude mouse <-> rat ES chimeras. Genes Cells 16, 397–405 (2011).

14. Kobayashi, T. et al. Generation of Rat Pancreas in Mouse by Interspecific Blastocyst Injection of Pluripotent Stem Cells. Cell 142, 787–799 (2010).

15. Goto, T. et al. Generation of pluripotent stem cell-derived mouse kidneys in Sall1-targeted anephric rats. Nat. Commun. 10, 451 (2019).

16. Usui, J. et al. Generation of Kidney from Pluripotent Stem Cells via Blastocyst Complementation. Am. J. Pathol. 180, 2417–2426 (2012).

17. Kitahara, A. et al. Generation of Lungs by Blastocyst Complementation in Apneumic Fgf10-Deficient Mice. Cell Rep. 31, 107626 (2020).

18. Mori, M. et al. Generation of functional lungs via conditional blastocyst complementation using pluripotent stem cells. Nat. Med. 25, 1691–1698 (2019).

19. Yamaguchi, T. et al. Interspecies organogenesis generates autologous functional islets. Nature 542, 191–196 (2017).

20. Chang, A. N. et al. Neural blastocyst complementation enables mouse forebrain organogenesis. Nature 563, 126–130 (2018).

21. Hamanaka, S. et al. Generation of Vascular Endothelial Cells and Hematopoietic Cells by Blastocyst Complementation. Stem Cell Reports 11, 988–997 (2018).

22. Ruiz-Estevez, M. et al. Liver development is restored by blastocyst complementation of HHEX knockout in mice and pigs. Stem Cell Res. Ther. 12, 292 (2021).

23. Kobayashi, T., Kato-Itoh, M. & Nakauchi, H. Targeted Organ Generation Using Mixl1-Inducible Mouse Pluripotent Stem Cells in Blastocyst Complementation. Stem Cells Dev. 24, 182–189 (2015).

24. Yamaguchi, T. et al. An interspecies barrier to tetraploid complementation and chimera formation. Sci. Rep. 8, 15289 (2018).

25. Liu, Y. & Hogan, B. L. Differential gene expression in the distal tip endoderm of the embryonic mouse lung. Gene Expr. Patterns 2, 229–233 (2002)

26. Okubo, T., Knoepfler, P. S., Eisenman, R. N. & Hogan, B. L. Nmyc plays an essential role during lung development as a dosage-sensitive regulator of progenitor cell proliferation and differentiation. Development 132, 1363–1374 (2005).

27. Porter, F. D. et al. Lhx2, a LIM homeobox gene, is required for eye, forebrain, and definitive erythrocyte development. Development 124, 2935–2944 (1997).

28. Shawlot, W. et al. Lim1 is required in both primitive streak-derived tissues and visceral endoderm for head formation in the mouse. Development 126, 4925–4932 (1999).

29. Tremblay, K. D., Hoodless, P. A., Bikoff, E. K. & Robertson, E. J. Formation of the definitive endoderm in mouse is a Smad2-dependent process. Development 127, 3079–3090 (2000).

30. Kanai-Azuma, M. et al. Depletion of definitive gut endoderm in Sox17-null mutant mice. Development 129, 2367–2379 (2002).

31. Oji, A. et al. CRISPR/Cas9 mediated genome editing in ES cells and its application for chimeric analysis in mice. Sci. Rep. 6, 31666 (2016).

32. Al Alam, D. et al. Evidence for the involvement of fibroblast growth factor 10 in lipofibroblast formation during embryonic lung development. Development 142, 4139–4150 (2015).

33. Okumura, H. et al. Contribution of rat embryonic stem cells to xenogeneic chimeras in blastocyst or 8-cell embryo injection and aggregation. Xenotransplantation 26, e12468 (2019).

34. Sui, P. et al. E3 ubiquitin ligase MDM2 acts through p53 to control respiratory progenitor cell number and lung size. Development 146, dev179820 (2019).

35. Franca, L. R., Ogawa, T., Avarbock, M. R., Brinster, R. L. & Russell, L. D. Germ cell genotype controls cell cycle during spermatogenesis in the rat. Biol. Reprod. 59, 1371–1377 (1998).

36. Rockich, B. E. et al. Spence, Sox9 plays multiple roles in the lung epithelium during branching morphogenesis. Proc. Natl. Acad. Sci. U.S.A. 110, E4456–E4464 (2013).

37. Moon, S. et al. Asxl1 exerts an antiproliferative effect on mouse lung maturation via epigenetic repression of the E2f1-Nmyc axis. Cell Death Dis. 9, 1118 (2018).

38. Bird, A. D. et al. cAMP Response Element Binding Protein Is Required for Differentiation of Respiratory Epithelium during Murine Development. PLOS ONE 6, e17843 (2011).

39. Kersbergen, A. et al. Lung morphogenesis is orchestrated through Grainyhead-like 2 (Grhl2) transcriptional programs. Dev. Biol. 443, 1–9 (2018).

40. O’Brien, K. B. et al. CARM1 is required for proper control of proliferation and differentiation of pulmonary epithelial cells. Development 137, 2147–2156 (2010).

41. Kalin, T. V. et al. Forkhead Box m1 transcription factor is required for perinatal lung function. Proc. Natl. Acad. Sci. U.S.A. 105, 19330–19335 (2008).

42. Kilkenny, C., Browne, W. J., Cuthill, I. C., Emerson, M. & Altman, D. G. Improving Bioscience Research Reporting: The ARRIVE Guidelines for Reporting Animal Research. PLOS Biol. 8, e1000412 (2010).

43. Choi, J. et al. Prolonged Mek1/2 suppression impairs the developmental potential of embryonic stem cells. Nature 548, 219–223 (2017).

44. Okada, Y. et al. Complementation of placental defects and embryonic lethality by trophoblast-specific lentiviral gene transfer. Nat. Biotechnol. 25, 233–237 (2007).

45. Kishimoto, Y. et al. A novel tissue specific alternative splicing variant mitigates phenotypes in Ets2 frame-shift mutant models. Sci. Rep. 11, 8297 (2021).

46. Isotani, A., Yamagata, K., Okabe, M. & Ikawa, M. Generation of Hprt-disrupted rat through mouse <-rat ES chimeras. Sci. Rep. 6, 24215 (2016).

