## supplementary figs and tables for "Generation of rat lungs by blastocyst complementation in Fgfr2b-deficient mouse model"

**This PDF file includes:**

Supplementary Figs. 1 to 8
Supplementary Tables 1 & 2

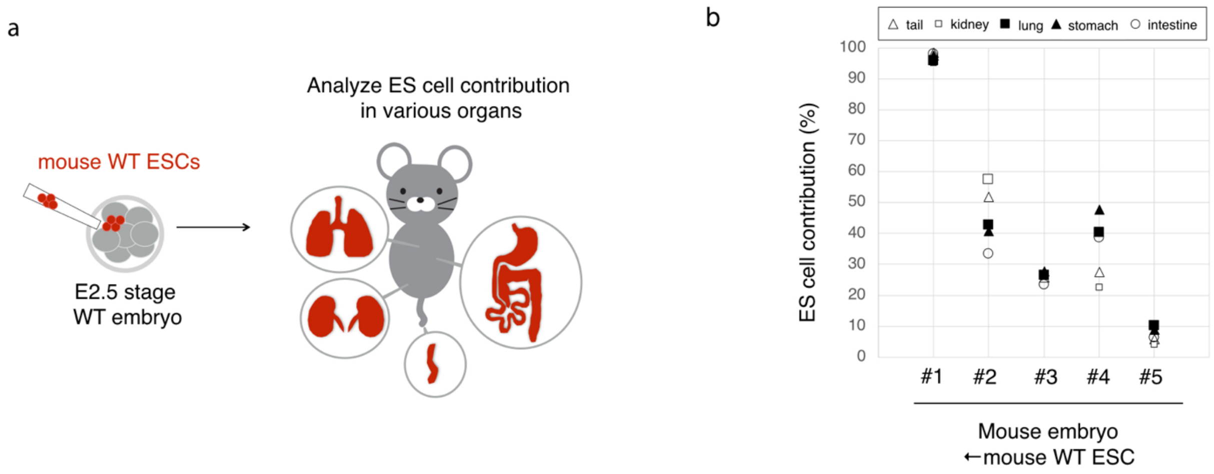


**Supplementary figure 1 Contribution of ESC-derived cells in mouse WT embryo ← mouse WT ESC chimeras in various tissues.**

**a** Red fluorescent protein (RFP)-expressing wild type (WT) embryonic stem cells (ESCs) were injected into the WT embryos at the E2.5 stage. The contribution of WT ESCs to various tissues was analyzed at E14.5.

**b** Flow cytometry results for RFP-expressing cells in the tail, kidney, lung, stomach, and intestine.

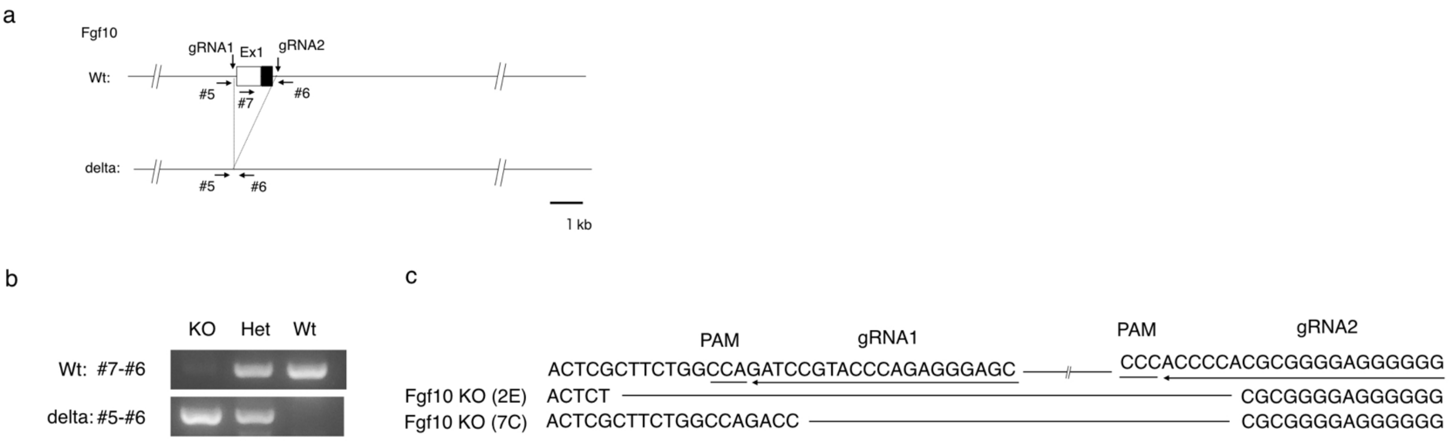


**Supplementary figure 2 Generation of Fgf10-KO ESC lines**

**a** Strategy of Fgf10 knockout (KO) model generation. Both gRNA1 and gRNA2 were used to remove exon1 of Fgf10.

**b** Genotype of Fgf10-KO, Fgf10 Heterozygous (Het), and Fgf10 WT ESCs.

**c** Mutation pattern of obtained Fgf10-KO ESC lines.

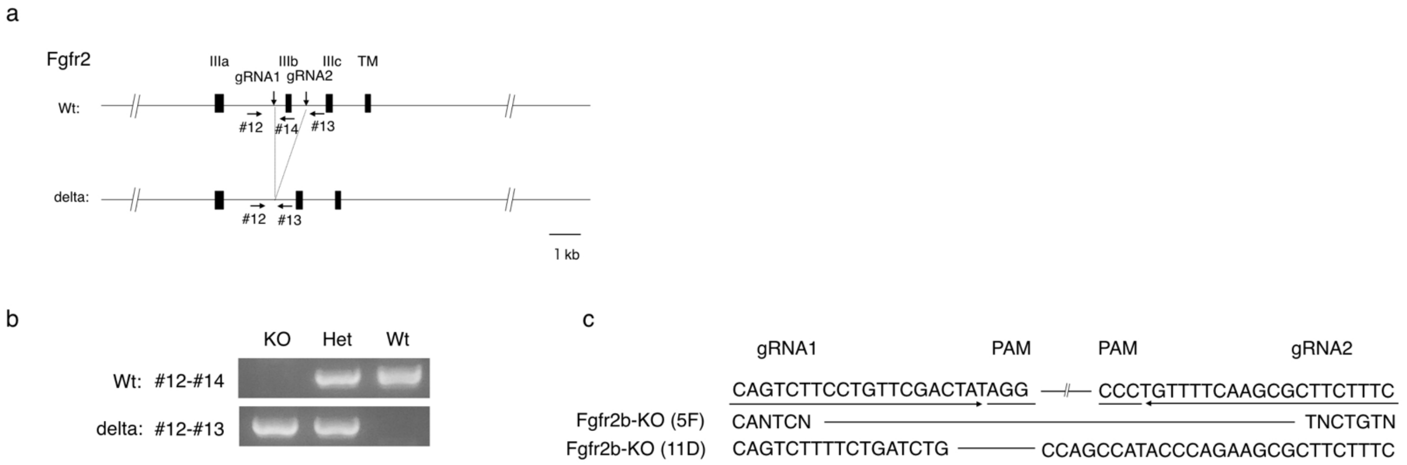


**Supplementary figure 3 Generation of Fgfr2b-KO ESC lines**

**a** Strategy of Fgfr2b-knockout (KO) model generation. Both gRNA1 and gRNA2 were used to remove the IIIb domain of Fgfr2.

**b** Genotype of Fgfr2b-KO, Fgfr2b heterozygous (Het), and Fgfr2b wild type (WT) ESCs.

**c** Mutation pattern of obtained Fgfr2b-KO ESC lines.

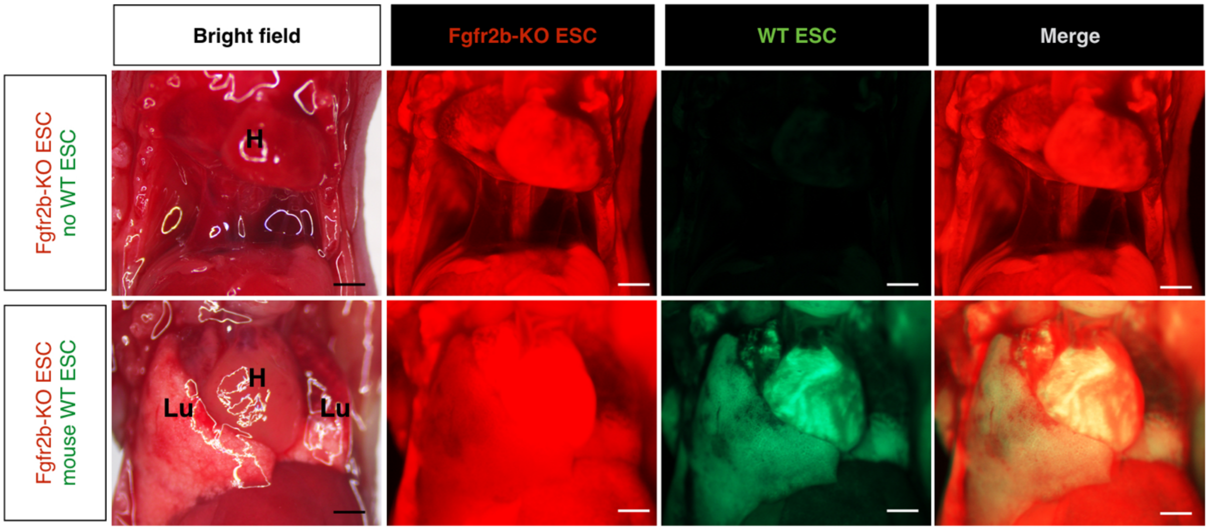


**Supplementary figure 4 RFP and GFP macroscopic images of heart and lung regions at P0.** Pups from Fgfr2b knockout (KO) embryonic stem cells (ESCs) only, and Fgfr2b-KO and wild type (WT) ESCs. Red fluorescent protein (RFP) is from Fgfr2b-KO ESC. Green fluorescent protein (GFP) is from mouse WT ESC. (H: heart, Lu: lung) Scale bars, 1 mm

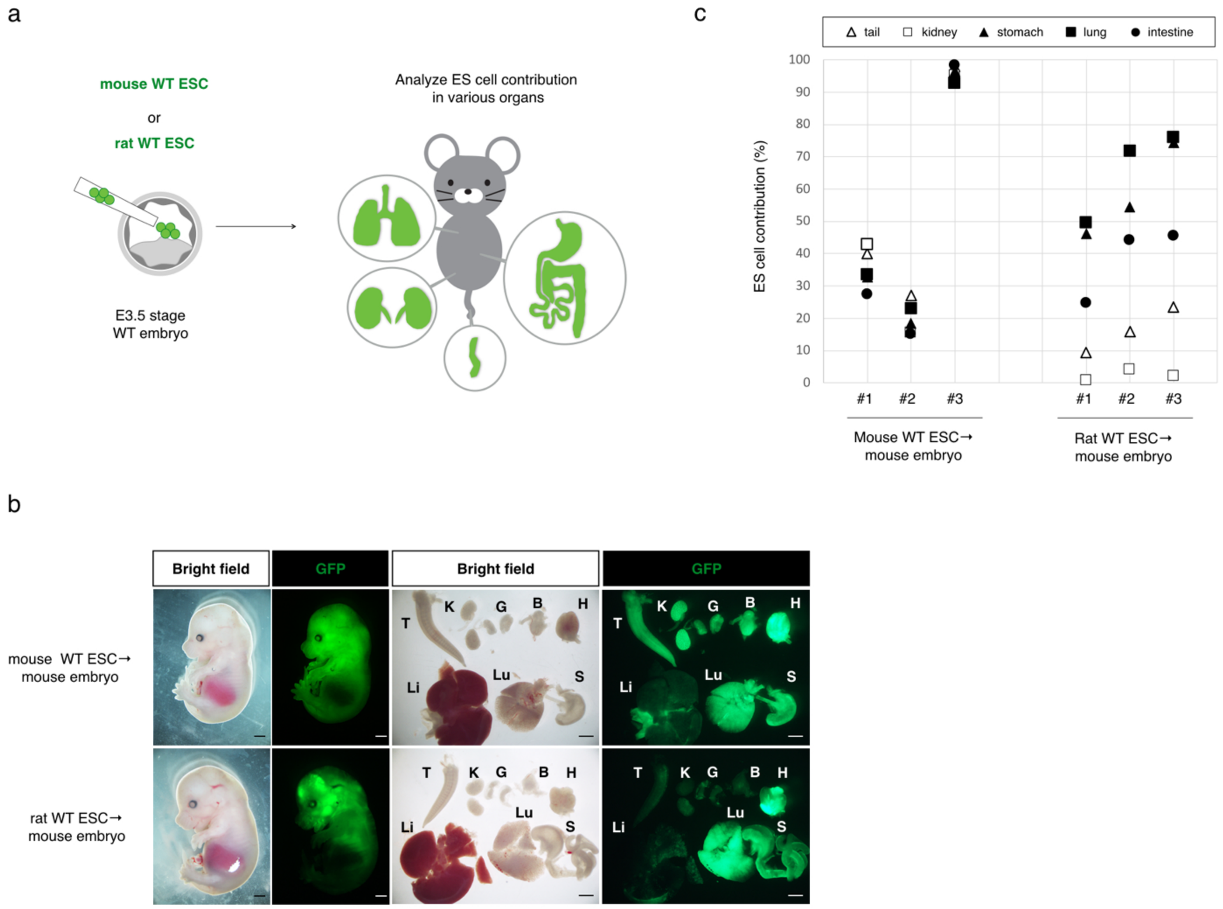


**Supplementary figure 5 Contribution of ESC-derived cells in various tissues from mouse WT embryo ← mouse WT ESC or rat WT ESC chimeras.**

**a** Green fluorescent protein (GFP)-expressing mouse wild type (WT) embryonic stem cells (ESCs) or rat WT ESCs were injected into WT embryos at the E3.5 stage. The contribution of WT ESCs to various tissues was analyzed at E14.5.

**b** Macroscopic and GFP image of embryo and tissues at E14.5. (T: tail, K: kidney, G: gonad, B: bladder, H: heart, Li: liver, Lu: lung, S: stomach)

**c** Flow cytometry results for GFP-expressing cells in the tail, kidney, lung, stomach or intestine. Note that rat cells are likely to contribute to the lung, but not to the kidney in mouse embryos.

a

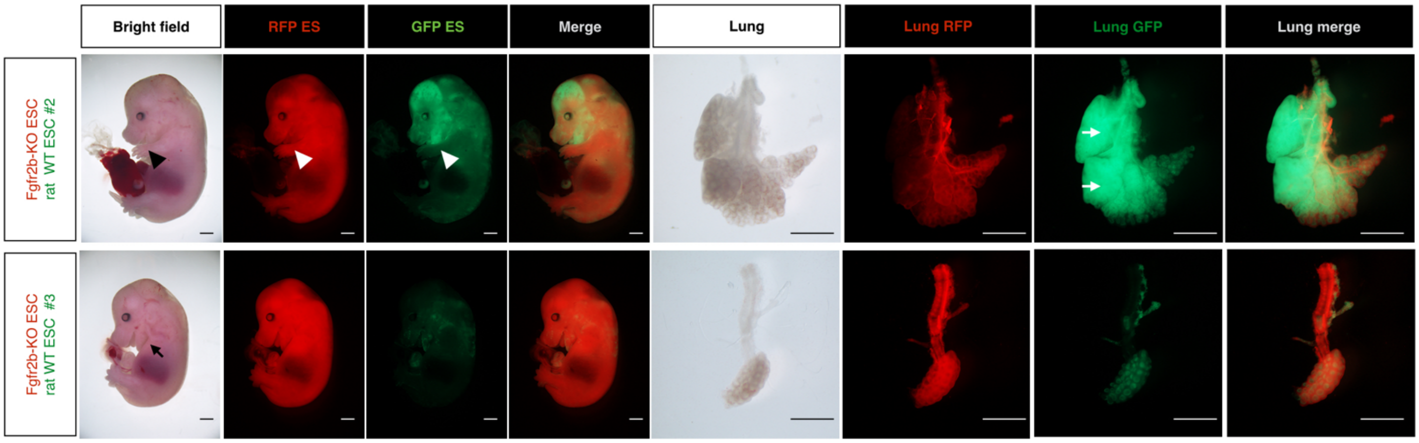


b

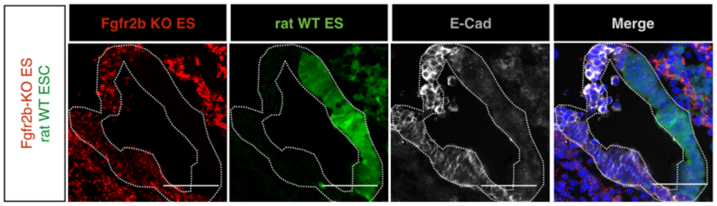


**Supplementary figure 6 Embryo and lung images from Fgfr2b-KO ESCs and rat WT ESCs chimeras.**

**a** Forelimb was observed in chimera #2 (black or white arrowheads). Parts of the lung lobes were mostly composed of rat cells (white arrows). When the rat cell contribution was low, forelimb was not observed in chimera #3 (black arrow) and the size of the lung was extremely small. Scale bars, 500 μm

**b** Immunostaining of E-Cad in lungs derived from Fgfr2b-knockout (KO) embryonic stem cells (ESCs) and rat wild type (WT) ESC chimeras. Note that part of the lung epithelial cells was composed of Fgfr2b-KO cells. Scale bars, 50 μm

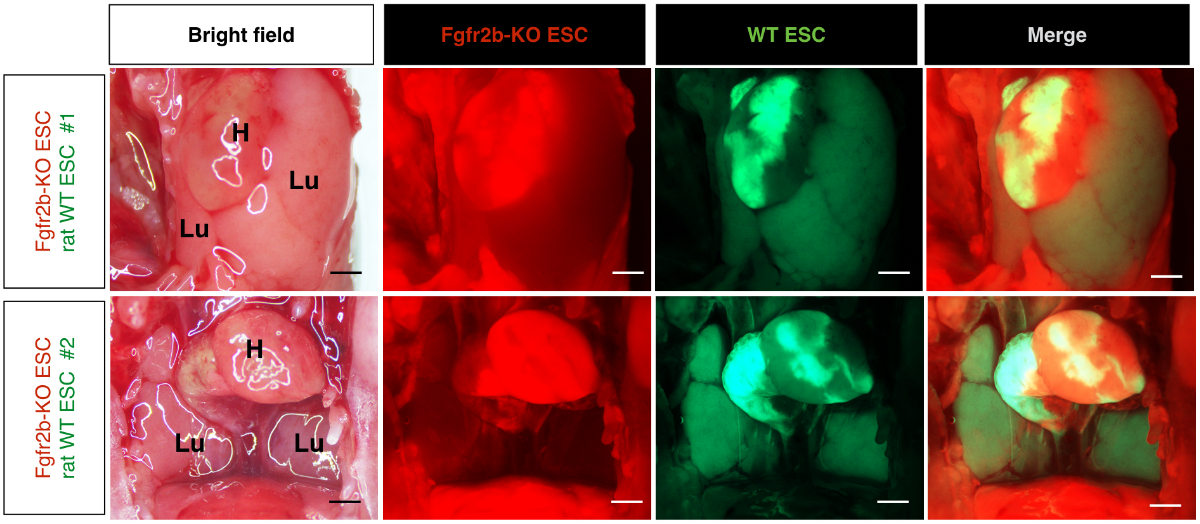


**Supplementary figure 7 RFP and GFP macroscopic images of heart and lung regions at P0.** Pups of Fgfr2b-knockout (KO) and rat embryonic stem cells (ESCs) (#1 and #2). Red fluorescent protein (RFP) is from Fgfr2b-KO ESC. Green fluorescent protein (GFP) is from rat wild type (WT) ESC. (H: heart, Lu: lung) Scale bars, 1 mm

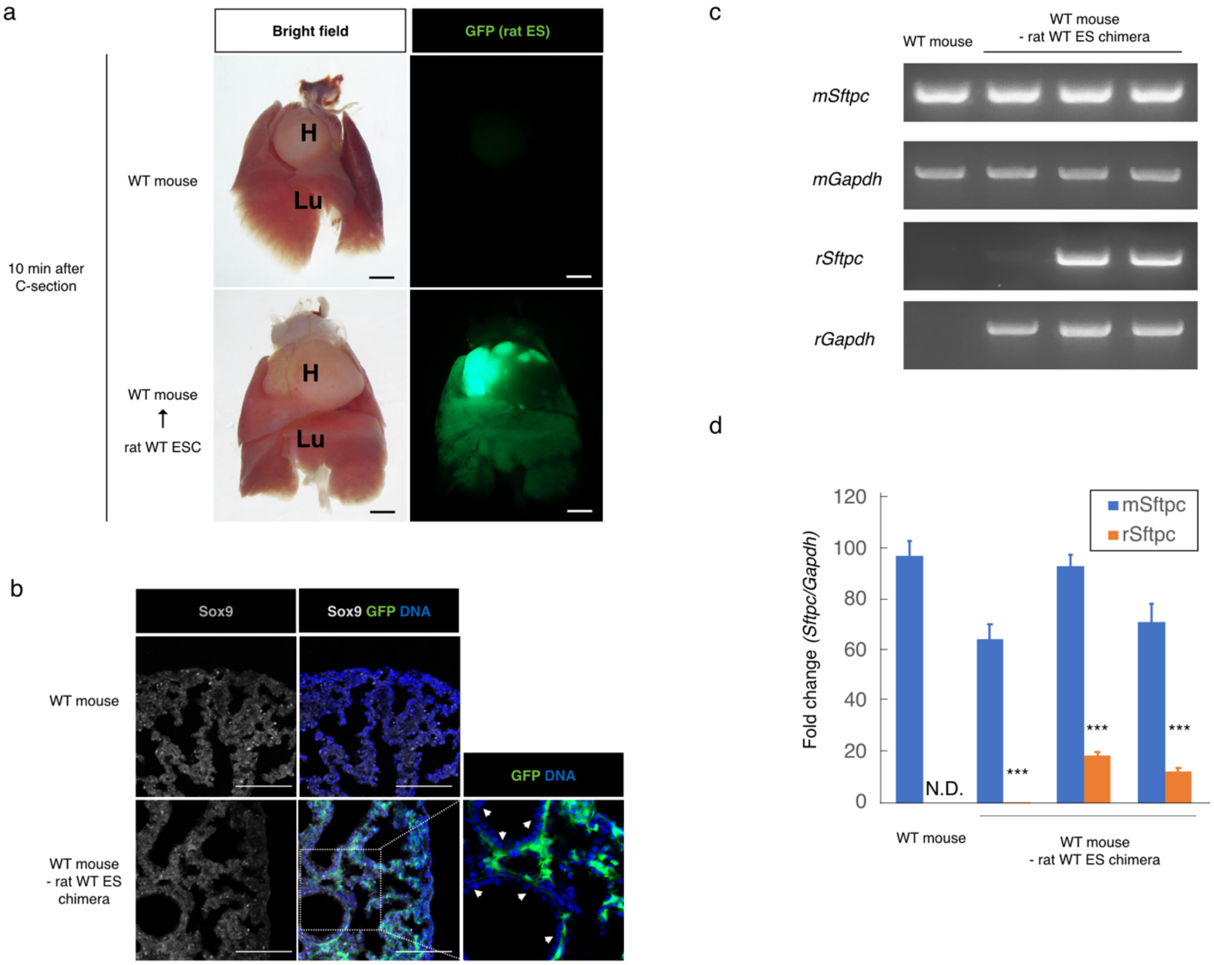


**Supplementary figure 8 Analysis of lungs in mouse WT embryo ← rat WT ESC chimera at P0**

**a** Macroscopic and green fluorescent protein (GFP) images of the lungs. (H: heart, Lu: lung)

**b** Immunostaining of Sox9 in lungs at P0 stage. Note absence of GFP-expressing cells from the epithelial tubules (white arrowhead). Scale bars, 100 µm

**c** RNA expression of mouse *Sftpc, Gapdh,* and rat *Sftpc, Gapdh* in the lungs at P0. RNA was extracted from one lung lobe for each sample.

**d** Quantitative real time PCR results of mouse *Sftpc* and rat *Sftpc*. Data were normalized to mouse or rat *Gapdh* expression levels. RNA was extracted from one lung lobe from each sample. N.D. not detected. ***: p<0.01

**Supplementary Table 1 Oligo DNA sequences.**

| ID | Oligo name | sequence |
|---|---|---|
| #1 | Fgf10-sgRNA1-F | caccGCTCCCTCTGGGTACGGATC |
| #2 | Fgf10-sgRNA1-R | aaacGATCCGTACCCAGAGGGAGC |
| #3 | Fgf10-sgRNA2-F | caccCCCCCCTCCCCGCGTGGGGT |
| #4 | Fgf10-sgRNA2-R | aaacACCCCACGCGGGGAGGGGGG |
| #5 | Fgf10 delta check F | GAGAAGGACCAGAAGGTGCC |
| #6 | Fgf10 delta check R | GGAATCAGGGTTGCAAGGGA |
| #7 | Fgf10-wt-checkF | GAGATGTCCGCTGGAGAAGG |
| #8 | Fgfr2b IIIb gRNA1 F | caccCAGTCTTCCTGTTCGACTAT |
| #9 | Fgfr2b IIIb gRNA1 R | aaacATAGTCGAACAGGAAGACTG |
| #10 | Fgfr2b IIIb gRNA2 F | caccGAAAGAAGCGCTTGAAAACA |
| #11 | Fgfr2b IIIb gRNA2 R | aaacTGTTTTCAAGCGCTTCTTTC |
| #12 | Fgfr2b IIIb delta check F | ACGGGATGGGGTAGATGGAA |
| #13 | Fgfr2b primer WT Rv | TGGATAGGATCCGGTGTGGA |
| #14 | FgfrIIIb-Wtcheck-Rv-mk2 | TCCCCGAGTGCTAGAACAGA |

**Supplementary Table 2 Primer sequences for RT-PCR.**

| Oligo name | sequence |
|---|---|
| RT-mGapdh F | CATTTGCAGTGGCAAAGTGGAG |
| RT-mGapdh R | CGTCAGATCCACGACGGAC |
| RT-mActa2 F | GCTGTGCTGTCCCTCTATGCC |
| RT-mActa2 R | GTACATGGTGGTACCCCCTGAC |
| RT-mE-Cad F | CGAGGAACCCTTTGAGGGGTCTCTTG |
| RT-mE-Cad R | CCTGGTCTTTGTTCTGGTTATCC |
| RT-mSftpc F | TCCCAGGAGCCAGTTCCGCATC |
| RT-mSftpc R | AGGTAGCGATGGTGTCTGCTCGC |
| RT-rGapdh F | CTTCTCTTGTGACAAAGTGGAC |
| RT-rGapdh R | ATGTCAGATCCACAACGGAT |
| RT-rSftpc F | AGTTTCGCATTCCCTGCTGC |
| RT-rSftpc R | GATGCCAGTGGAGCCAATAGAG |
